# Interactions among mTORC, AMPK, and SIRT: A Computational Model for Cell Energy Balance and Metabolism

**DOI:** 10.1101/2020.10.07.330308

**Authors:** Mehrshad Sadria, Anita T. Layton

## Abstract

Key proteins such as mTORC, AMPK, and sirtuins are known to play an essential role in the management of metabolic stress and ageing mechanisms. An impairment in these mechanisms is commonly associated with cellular ageing and degenerative diseases. To understand the complex interactions of ageing□related signalling pathways and environmental signals, and the impacts on lifespan and health-span, we developed a computational model of ageing signalling pathways. The model includes (i) the insulin/IGF-1 pathway, which couples energy and nutrient abundance to the execution of cell growth and division, (ii) mTORC1 and amino acid sensors, (iii) the Preiss-Handler and salvage pathways, which regulate the metabolism of NAD+ and the NAD+-consuming factor SIRT1, (iv) the energy sensor AMPK, and (v) transcription factors FOXO and PGC-1α. Key findings include the clinically important role of PRAS40, sestrin2, and diet in the treatment of cancers and other diseases, and a potential link between SIRT1-activating compounds and premature autophagy. The model can be used as an essential component to simulate gene manipulation, therapies (e.g., rapamycin and wortmannin), calorie restrictions, and chronic stress, and to assess their functional implications on longevity and ageing□related diseases.

**Author Summary:** In cellular ageing, mitochondrial function declines over time, which affects normal mechanisms of cells and organisms and leads to myriad of degenerative diseases and other health problems. To investigate the mechanisms that affect the ageing process, we focus on pathways that play a key role in the management of metabolic stress: the mTORC, AMPK, and sirtuins pathways. Our goal is to understand the complex interactions of ageing and metabolism related signalling pathways and environmental signals, and the impacts on lifespan and health-span. To accomplish that goal, we developed a computational model of signalling pathways related to ageing and metabolism. By conducting model simulations, we have unraveled the clinically important role of PRAS40, sestrin2, and diet in the treatment of cancers and other diseases, and a double-edged sword effect of SIRT1-activating compounds in their use as a health remedy. We view this model as an essential step towards a tool for studying metabolism, longevity, and ageing-related diseases. By extending the present model as appropriate, we can simulate gene manipulation, therapies (e.g., rapamycin and wortmannin), calorie restrictions, and chronic stress,.

## INTRODUCTION

Recent medical advances have drastically increased life expectancy. By 2050 the world’s population aged 60 years and older is expected to total 2 billion, up from 900 million in 2015.(1) However, the prevalence of age-related diseases such as cancer, diabetes, and neurodegenerative and cardiovascular diseases has risen accordingly. Ageing is a multifactorial process characterized by a gradual decline of physiological functions. A series of mechanisms are involved at the molecular, cellular, and tissue levels, which include deregulated autophagy, mitochondrial dysfunction, telomere shortening, oxidative stress, systemic inflammation, and metabolism dysfunction.(2) The deregulation of these pathways gives rise to cellular senescence, which contributes to ageing phenotype and, eventually, age-related diseases.

Despite intense research, the molecular basis of ageing has remained incompletely understood. That challenges may be attributed to the multitude of molecular and cellular processes involved. In particular, the insulin/IGF-1 (Insulin-like Growth Factor 1) signaling pathway are known to regulate lifespan, and is coupled to the mechanistic target of rapamycin (mTOR) pathway. mTOR is a highly conserved serine/threonine protein kinase with two distinct complexes, mTORC1 and mTORC2. mTOR controls cell growth, proliferation, motility and survival, protein and lipid synthesis, glucose metabolism, mitochondrial function and transcription, in response to nutrient and hormonal signals. Inhibition of this nutrient response pathway extends lifespan in model organism and ameliorates age-related pathologies. The presence of nutrients and growth factors increases mTORC1 activity and reduces autophagy initiation, whereas the caloric restriction or inactivation of mTORC1 increases autophagy contributing to cellular longevity. Insulin/IGF-1 activates mTORC1 by AKT (a.k.a. protein kinase B), while mTORC1 inhibits insulin/IGF-1 through S6K by inhibiting insulin receptor substrate (IRS).

Also important in the ageing process is the sirtuin family, which consists of highly conserved protein deacetylases found nearly in all organisms studied. In mammals, seven silent information regulator (SIRT) proteins (SIRT1-7) exist, with SIRT1 being the most extensively studied in the context of ageing and is known as one of the main mediators in calorie restriction. SIRT1 senses changes in intracellular nicotinamide adenine dinucleotide (NAD+) levels, and uses this information to adapt the cellular energy output. SIRT1 also enhances DNA repair, cell survival, mitochondrial function and reduces ageing inflammatory/immune responses.(3)

Recent high-throughput genomic and proteomic technologies have generated a wealth of ageing-related data. Nonetheless, some of the molecular mechanisms that mediate key ageing effects have yet to be elucidated. The difficulty lies in the complexity of ageing: Not only are a large number of genes involved in the ageing process, many with competing roles, but their interactions are complex and often incompletely characterized. Indeed, due to the multiple feedback loops and regulatory mechanisms, it is challenging to understand the biological consequences of gene-expression changes. A promising methodology for interpreting data and untangling the interactions among signaling pathways is computational biology. One such approach is to describe regulatory interactions using ordinary differential equations (ODEs), which relate changes in the expressions of model variables to other quantities. The insulin/IGF-1 pathway has been the subject of modeling and analysis by a series of previous studies.(4–11)

Unlike the insulin/IGF-1 and mTOR pathways, theoretical effort in modeling the arguably equally important regulators NAD+ and SIRT1 is limited (an exception is (12)). Thus, the goal of this study is to develop a state-of-the-art computational model that couples these and other critical signaling pathways in growth, ageing, metabolism, and disease in mammals. To achieve that goal, we present a comprehensive model that includes (i) the insulin/IGF-1 pathway, which couples energy and nutrient abundance to the execution of cell growth and division, (ii) mTORC1 and the amino acid (AA) sensors, (iii) the Preiss-Handler and salvage pathways, which regulate the metabolism of NAD+ and the NAD+-consuming factor SIRT1, (iv) the energy sensor adenosine monophosphate-activated protein kinase (AMPK), and (v) transcription factors forkhead box O (FOXO) and peroxisome proliferator-activated receptor gamma coactivator 1-α (PGC-1α), the overexpression or mutation of which affects lifespan. We apply the model to investigate the synergy among regulators of nutrients, energy, metabolism, and autophagy, and to identify novel therapeutic targets. The model can be used to aid in the interpretation of genomic and proteomic data, and to provide an integrated understanding of the mechanisms that lead the cell to senescence and how this process contributes to ageing and age-related diseases.

## RESULTS

### PRAS40 substantially lowers mTORC1 level under low insulin conditions

Locating at the crossroad of the insulin/IGF-1 pathway, the proline-rich AKT substrate of 40 kDa (PRAS40) is phosphorylated by growth factors or other stimuli, and in turn regulates the activation of these signaling pathways. PRAS40 plays an important role in metabolic disorders and multiple cancers, and is known to be an insulin-regulated inhibitor of mTORC1.(13) To assess how that regulation is altered under pharmacological interventions, we compare model predictions with and without PRAS40 inhibition of mTORC1. In these simulations, the model is initialized at the fasting state with a low insulin level (10% baseline). At t=40 min, insulin level is returned to baseline.

With sufficient insulin and no drug treatment (i.e., control; administration of rapamycin will be considered below), PRAS40 inhibition of mTORC1 has only a minor impact on mTORC1, lowering its steady-state phosphorylated level by 19% (compare Fig. 1a, left and right panels, black curves, t > 40 min). The effects of PRAS40 on other proteins are similarly minor (Figs. 1 and 2; Fig. S2, Supplemental Materials).

**Figure 1.**
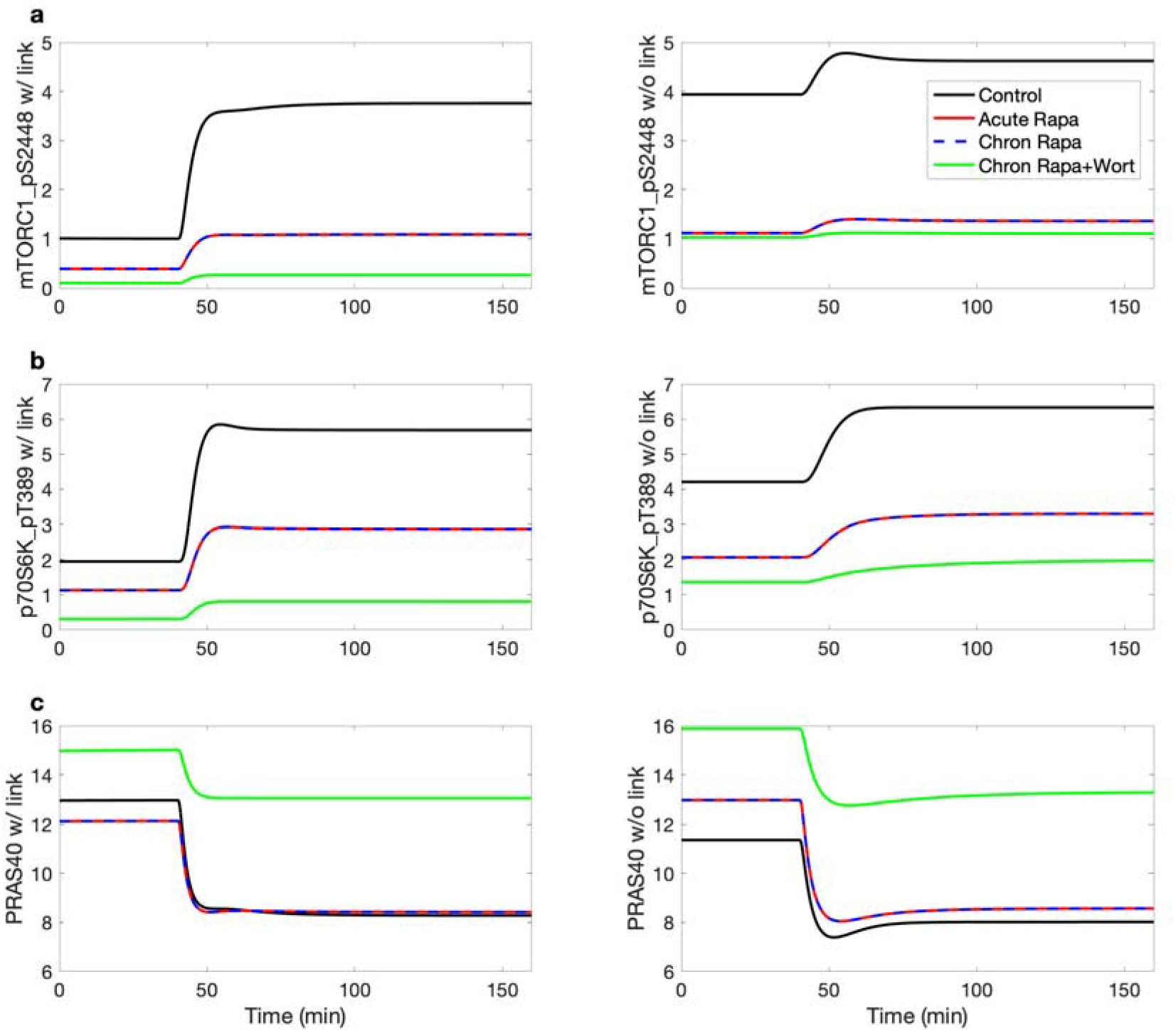
Effects of insulin, rapamycin, and wortmannin on key proteins and their interactions. Insulin was lowered to 10% of its baseline level for the initial 40 min of the simulation, and subsequently returned to baseline level. Simulations are conducted for control, acute and chronic administration of rapamycin, chronic administration of rapamycin with wortmannin. The inhibition of PRAS40 of mTORC1 is represented in the left panels but not the right ones. Model predicts that PRAS40 substantially lowers mTORC1 level under low insulin conditions. That effect is the most prominent under chronic administration of rapamycin and wortmannin.

**Figure 2.**
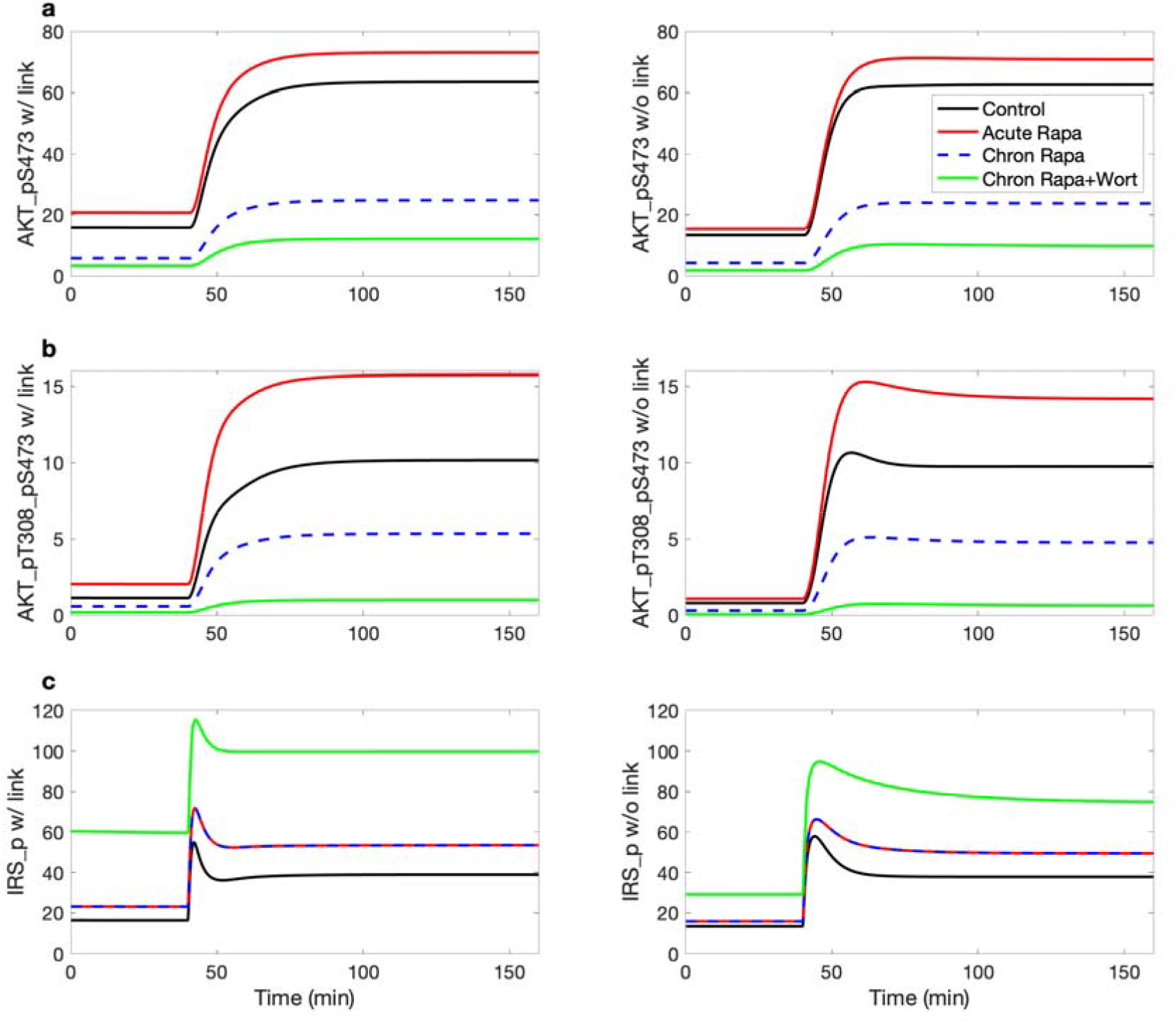
Effects of insulin, rapamycin, and wortmannin on key proteins and their interactions. Notations are analogous to Fig. 1.

At low insulin level, PRAS40 inhibition of mTORC1 results in a substantially lower phosphorylated mTORC1 level (1.00 versus 3.94; compare Fig. 1a panels, black curves, t < 40 min). A lower insulin level decreases AKT_pT308_S473 (Fig. 2b) and thus phosphorylated mTORC1; both changes reduce the phosphorylation rate of PRAS40 (Fig. 1c). The resulting higher unphosphorylated PRAS40 level reduces mTORC1 phosphorylation rate.

### Rapamycin alone does not significantly alter PRAS40 levels

Acute rapamycin administration lowers phosphorylated mTORC1 level and decreases the phosphorylation of PRAS40 to PRAS40_pS183, which, taken in isolation, would increase dephosphorylated PRAS40. However, the lower phosphorylated mTORC1 level also decreases p70S6K_pT389, increases phosphorylated IRS and phosphorylated PI3K_PDK1, and eventually, increases AKT_pT308 and AKT_pT308_pS473, both of which increase the phosphorylation of PRAS40 to PRAS40_pT246. The competing effects on PRAS40 phosphorylation results in negligible change in unphosphorylated PRAS40 level.

With chronic rapamycin administration, the lower phosphorylated mTORC1 slows the phosphorylation of PRAS40 to PRAS40_pS183. The lower phosphorylated mTORC2 level slows the phosphorylation of AKT to AKT_pS473, reducing both AKT_pS473 and the downstream AKT_pT308_pS473 (Fig. 2(a,b)). Taken in isolation, these effects, together with the reduced phosphorylated mTORC1, would slow the phosphorylation of PRAS40 and increase dephosphorylated PRAS40 level. But in a competing effect, the lower phosphorylated mTORC1 level increases AKT_pT308, which increases the phosphorylation of PRAS40 to PRAS40_pT246. These competing effects together yield negligible change in unphosphorylated PRAS40 level (Fig. 1c).

### The addition of wortmannin significantly elevates PRAS40 and further suppresses mTORC1

A noticeable effect can be observed in PRAS40 when wortmannin or MK-2206 is combined with chronic rapamycin administration. The phosphorylated PI3K_PDK1 level is lowered, which decreases AKT_pT308 and AKT_pT308_pS473, slowing the phosphorylation of PRAS40 to PRAS40_pT246. Together with reduced phosphorylated mTORC1, this maneuver substantially increases unphosphorylated PRAS40 at baseline insulin by 55% (Fig. 1c, left). The elevated PRAS40 substantially suppresses mTORC1_pS2448 (Fig. 1a, left). Analogous effects are also obtained for MK-2206, an allosteric inhibitor of AKT.

### Optimizing rapamycin dosage to maintain insulin sensitivity while preserving mTORC1 inhibition

Chronic administration of rapamycin inhibits mTORC2 and impair insulin sensitivity by reducing AKT_pT308_pS473, which is essential in the translocation of GLUT4.(14, 15) We aim to determine an optimal rapamycin dosage that attenuates the detrimental effect on insulin sensitivity while preserving mTORC1 inhibition. We simulate the glucose tolerance test under four conditions: control, acute rapamycin administration, chronic rapamycin administration, and chronic rapamycin and wortmannin administration (parameters as described in previous simulations). For chronic rapamycin, we considered mTORC2 inhibition at 75% (baseline), but also at 65%, 50%, and 25%, with mTORC1 inhibition fixed at 75%.

The predicted plasma glucose time-course profiles are shown in Fig. 3, together with experiment data obtained for vehicle.(16) Acute rapamycin usage inhibits mTORC1 (but not mTORC2), which suppresses p70S6K_pT389, increases phosphorylated IRS, AKT_pT308 and AKT_pT308_pS473, thereby improving insulin sensitivity. In contrast, chronic rapamycin usage inhibits mTORC2 as well. That lowers AKT_pS473 and AKT_pT308_pS473, and, at sufficiently high dosage (>50% mTORC2 inhibition), leads to impaired glucose tolerance. If wortmannin is added, AKT_pT308_pS473 is further suppressed, resulting in a sustained elevated plasma glucose level.

**Figure 3.**
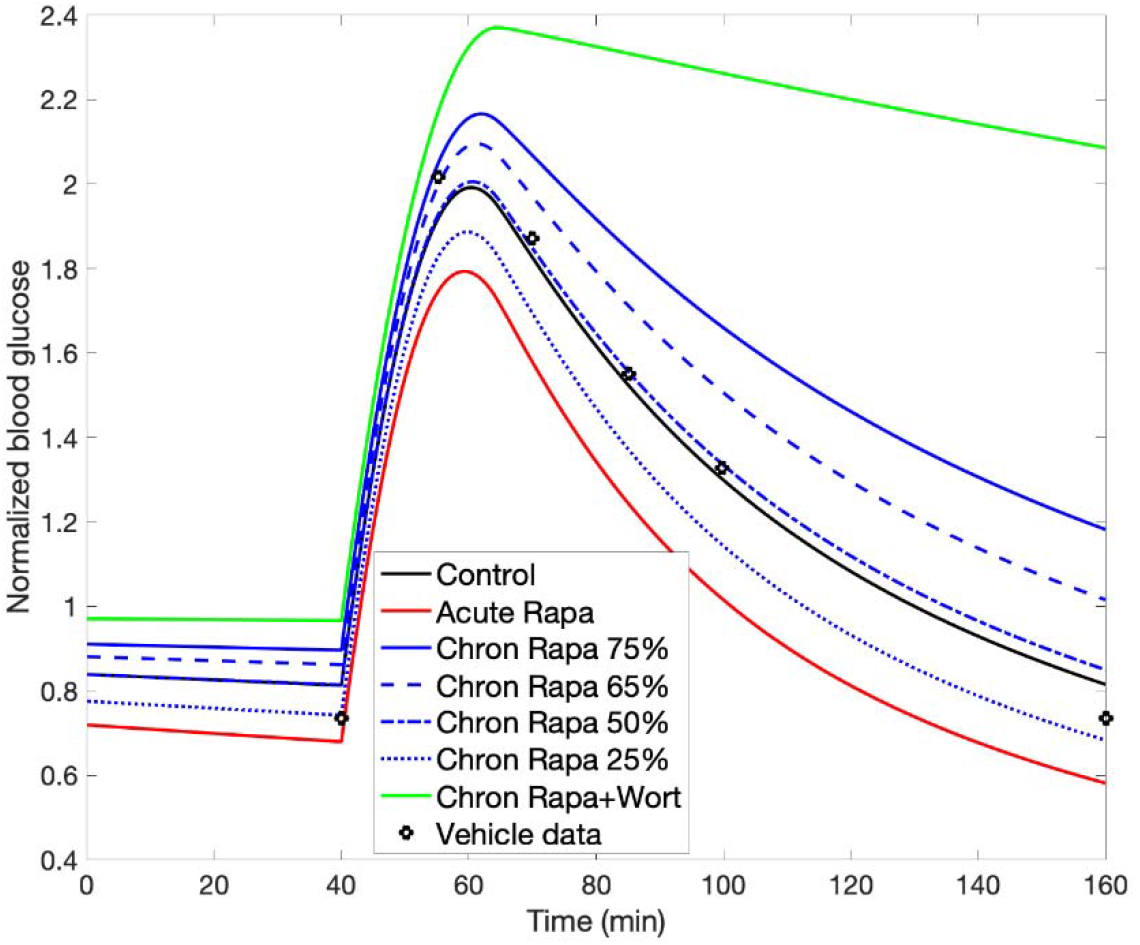
Effect of rapamycin on glucose tolerance. Glucose tolerance test begins at t = 40 min. Normalized plasma glucose profiles are obtained for acute rapamycin administration, and for chronic rapamycin administration at different dosages, with and without wortmannin. Vehicle data from (16).

Findings in PC3 cells suggest that differing rapamycin dosages may yield near maximal mTORC1 inhibition with a range of mTORC2 inhibition levels.(17) Model simulations suggest that lowering rapamycin-induced mTORC2 inhibition from 75% (baseline) to 50% restores insulin sensitivity to control level, consistent with intermittent administration of rapamycin.(18, 19) If it is possible to further reduce mTORC2 inhibition to 25% while preserving mTORC1 inhibition at 75%, one even achieves an improvement in insulin sensitivity, due to the beneficial effect of mTORC1 inhibition overriding the impairment arising from the (attenuated) mTORC2 inhibition.

### Regulation of mTORC1 by arginine and leucine

During AA starvation, lysosomal AAs (leucine in particular) facilitate mTORC1 activation. The transfer of lysosomal leucine to cytoplasm is mediated by SLC38A9, an arginine-regulated AA transporter. Thus, leucine efflux depends on the concentrations of both lysosomal leucine and arginine (Fig. 4(a,b)). To assess the role of arginine in the regulation of mTORC1 and autophagy during AA depletion, we conduct simulations in which cytoplasmic AA progressively decreases during the first 3 hours; afterwards, lysosomal AAs are released.(20) We simulate high, medium, low, and zero arginine levels by varying the rate at which cytoplasmic AA increases due to lysosomal leucine efflux. In the presence of insulin, AAs activate mTORC1, which inhibits autophagy by phosphorylating ULK1. (21) Upon AA depletion, mTORC1 activation on the lysosomal surface is no longer maintained (Fig. 4c); a similar trend is observed for the mTORC1 readout p70S6K (Fig. 4d). Consequently, ULK1 Ser757 is rapidly dephosphorylated (Fig. 4e),(21) resulting in activation of the ULK1 kinase and concomitant autophagy induction. When cytoplasmic AAs are sufficiently depleted, leucine is transported out of the lysosome by arginine-stimulated SLC38A9, activating mTORC1, phosphorylating p70S6K and ULK1, and suppressing autophagy (Fig. 4(c,d)). Additional results can be found in Fig. S3.

**Figure 4.**
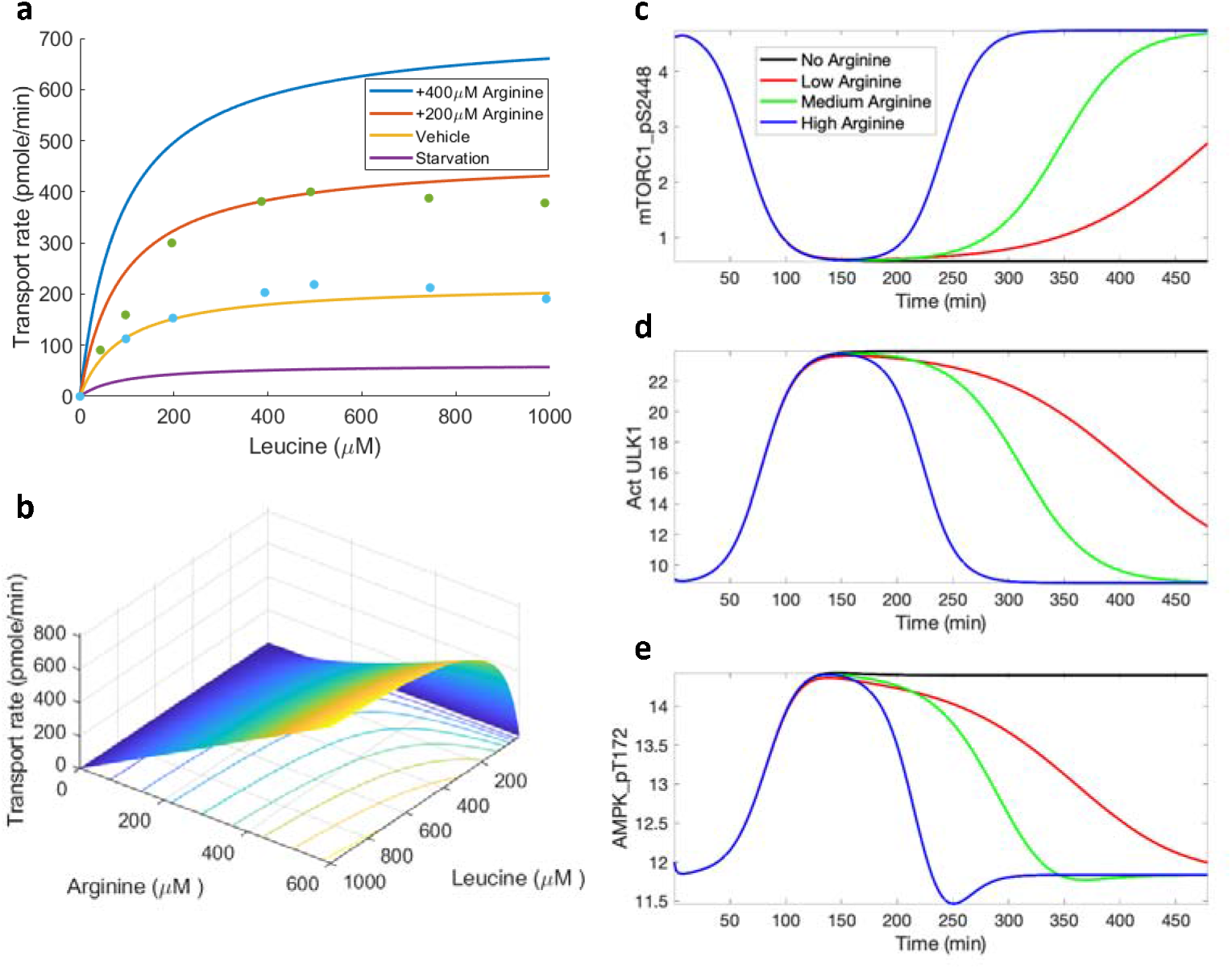
Leucine transport rate as a function of leucine and arginine concentrations, and its effect on mTORC1 reactivation. Panel ***a***, leucine transport rate is computed with arginine concentration taken to be 50 μM (“Starvation”), 175 μM (“Vehicle”), 375 μM (“+200 μM Arginine”), 575 μM (“+400 μM Arginine”). Data taken from (48). Panel ***b***, leucine transport rate shown for the full range of leucine and arginine concentrations. Predicted levels of mTORC1 pS2448 (***c***), activated ULK1 (***d***), and AMPK_pT172 (***e***) under amino acid depletion, obtained for differing arginine levels.

### Regulation of mTORC1 by sestrin2 and leucine

Sestrin has been found to induce autophagy during diverse environmental stresses that provoke mitochondrial dysfunction,(22) through AMPK activation and mTORC1 inhibition. Specifically, after leucine binds to sestrin2, the resulting complex activates AMPK. Thus, the model assumes that AMPK phosphorylation rate is proportional to the concentration product [Leucine][Sestrin].(23) How does the interaction between sestrin2 and AAs affect the dynamics of AMPK activation, mTORC1 inhibition, and autophagy stimulation? To answer that question, we conduct simulations in which AA levels (including leucine) are initially set to 10% of baseline values, then subsequently increased to baseline levels. Simulations are conducted for high, medium, and low sestrin2 levels.

The regulation of mTORC1 activity by leucine and sestrin2 is shown in Fig. 5(a,b), by evaluating Eq. 4 for given AA and TSC1_TSC2 complex levels. Consider the system with a typical lysosomal leucine concentration. At a sufficiently low sestrin2 concentration, GATOR2 is activated, GATOR1 is inhibited, and mTORC1 activation rate is maximized. Conversely, at sufficiently high sestrin2 concentration, mTORC1 is rapidly dephosphorylated. Also, for a fixed sestrin2 concentration, increasing the concentration of leucine raises the phosphorylation rate of mTORC1. Now for a typical sestrin2 concentration, Fig. 5a exhibits the Michaelis-Menten-like dependence of mTORC1 activity on leucine.

**Figure 5.**
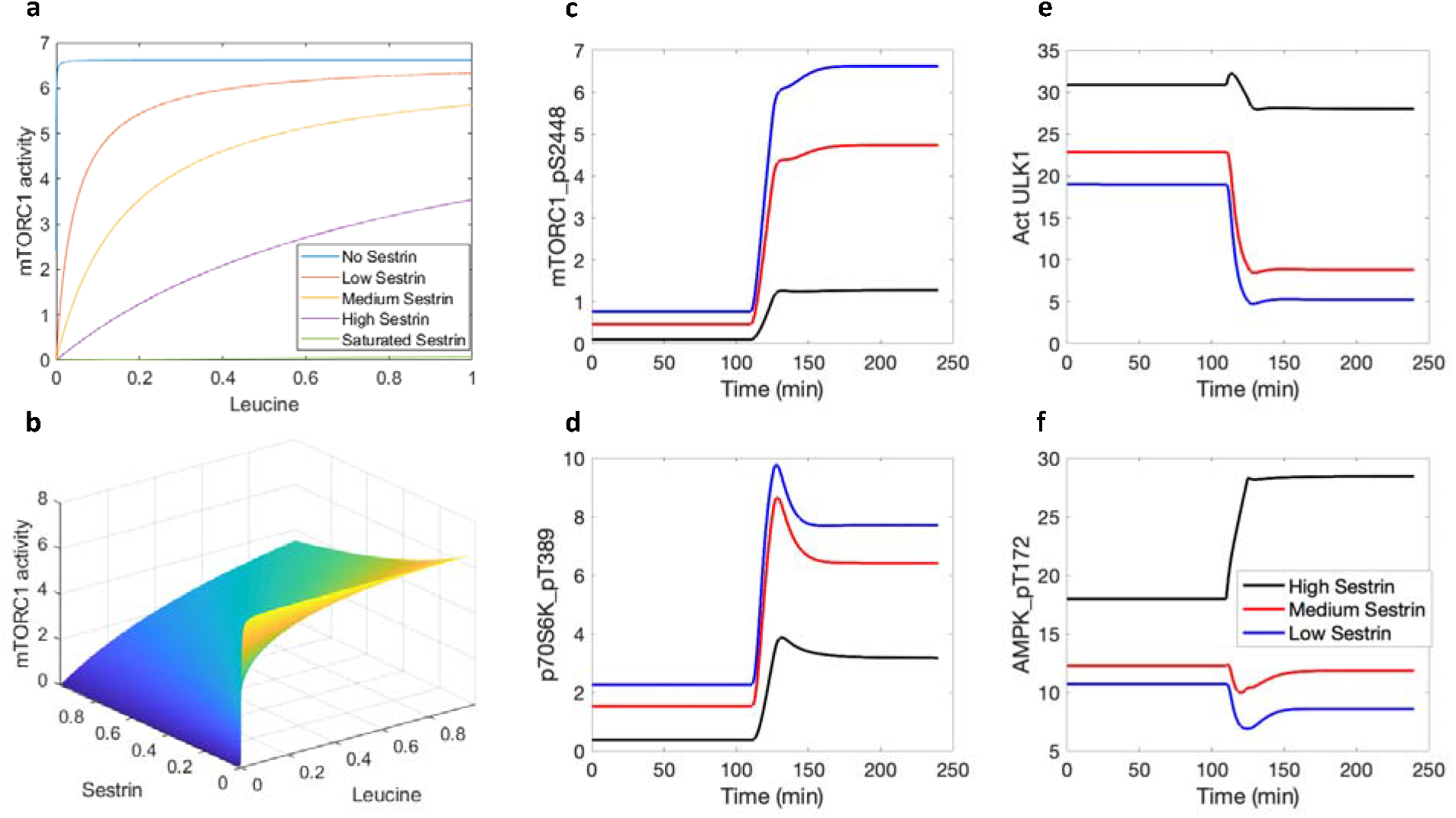
mTORC1 activity as a function of normalized leucine and sestrin concentrations, and model response to protein depletion and restoration. Panel ***a***, mTORC1 activity for differing sestrin levels. Panel ***b***, mTORC1 activity for the full range of leucine and sestrin concentrations. Predicted levels of mTORC1_pS2448 (***c***), p70S6K_pT389 (***d***), activated ULK1 (***e***), and AMPK_pT172 (***f***) under amino acid depletion (t < 2 h) and restoration (t > 2h), obtained for differing sestrin levels.

Key model variables are exhibited in Figs. 5 and S4. Time profiles of mTORC1_S2448 and p70S6K_pT389 approximate that of AAs, whereas activated ULK1 exhibits the opposite trend. As sestrin2 concentration decreases, the activating effect of AAs on mTORC1 and p70S6K is enhanced (Fig. 5(c,d)), as is their inhibition of ULK1 and autophagy (Fig. 5e). AAs have competing effects on the phosphorylation of AMPK. Leucine and sestrin2 together activate AMPK directly. In addition, through mTORC1 and its inhibition of IRS_p, AAs inhibit AMPK. The model predicts that the latter (inhibition) dominates, resulting in the activation of AMPK when proteins are depleted, consistent with experimental observations.(24) The inhibition of AMPK by AAs is modulated by sestrin2 (Fig. 5f).

### STACs and their anti-ageing effects

Resveratrol, a SIRT1 activator, mimics the anti-ageing effects of calorie restriction in lower organisms and mice.(25) Other SIRT1-activating compounds (STACs) have been identified that are structurally unrelated to and more potent than resveratrol.(25) As shown in Fig. 6a, the STACs increase SIRT activities to different degrees. SIRT regulates transcription by deacetylating transcription factors FOXO.(26) Model predicts that all STACs increase FOXO deacetylation but to significantly different degrees (Fig. 6b). At baseline K_m_ of [FOXO] = 141μm, resveratrol increases FOXO deacetylation rate by a moderate degree (11%). Other STACs are more effective, with the largest increase obtained for SRT1720 (54%).

**Figure 6.**
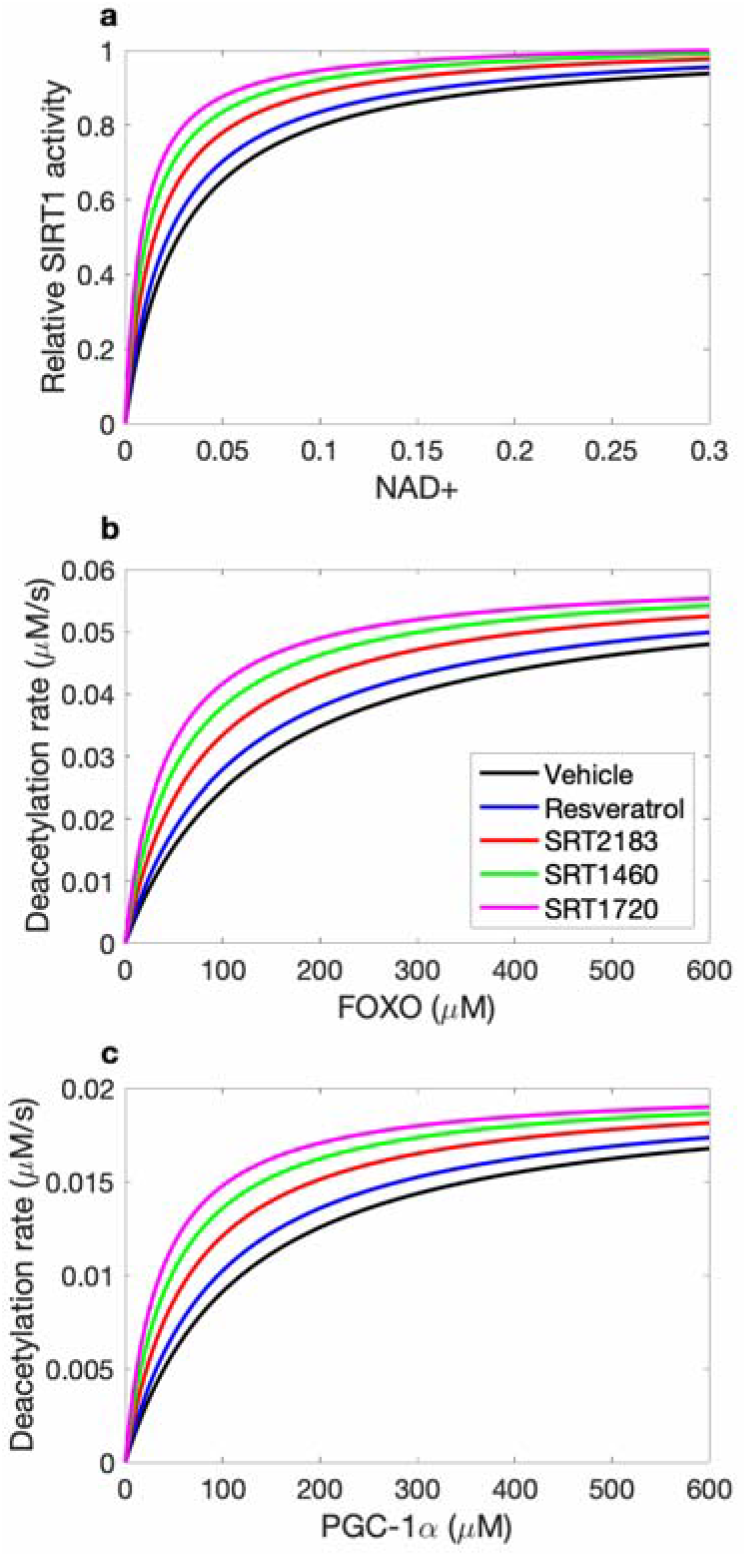
STACs activate SIRT1 (normalized values shown) by lowering the Michaelis-Menten constant for NAD+ (panel ***a***). The deacetylation rates of FOXO and PGC-1α are subsequently affected (panels ***b*** and ***c***).

STACS activate AMPK and SIRT1, leading to the deacetylation of peroxisome proliferator-activated receptor-γ coactivator 1-α (PGC-1α). PGC-1α activation improves mitochondrial biogenesis. At baseline PGC-1α K_m_ level, resveratrol increases PGC-1α deacetylation rate by 12%. Larger improvements are obtained for other STACs (Fig. 6c).

The effects of these compounds on key protein activities are shown in Fig. 7. Consistent with results in Fig. 6a, STACs raise SIRT1 levels, by as much as 32% (SRT1720; Fig. 7a). SIRT1 in turn activates AMPK, although the SIRT1-induced increase in AMPK_pT172 is relatively small compared to SIRT1 (Fig. 7b).

**Figure 7.**
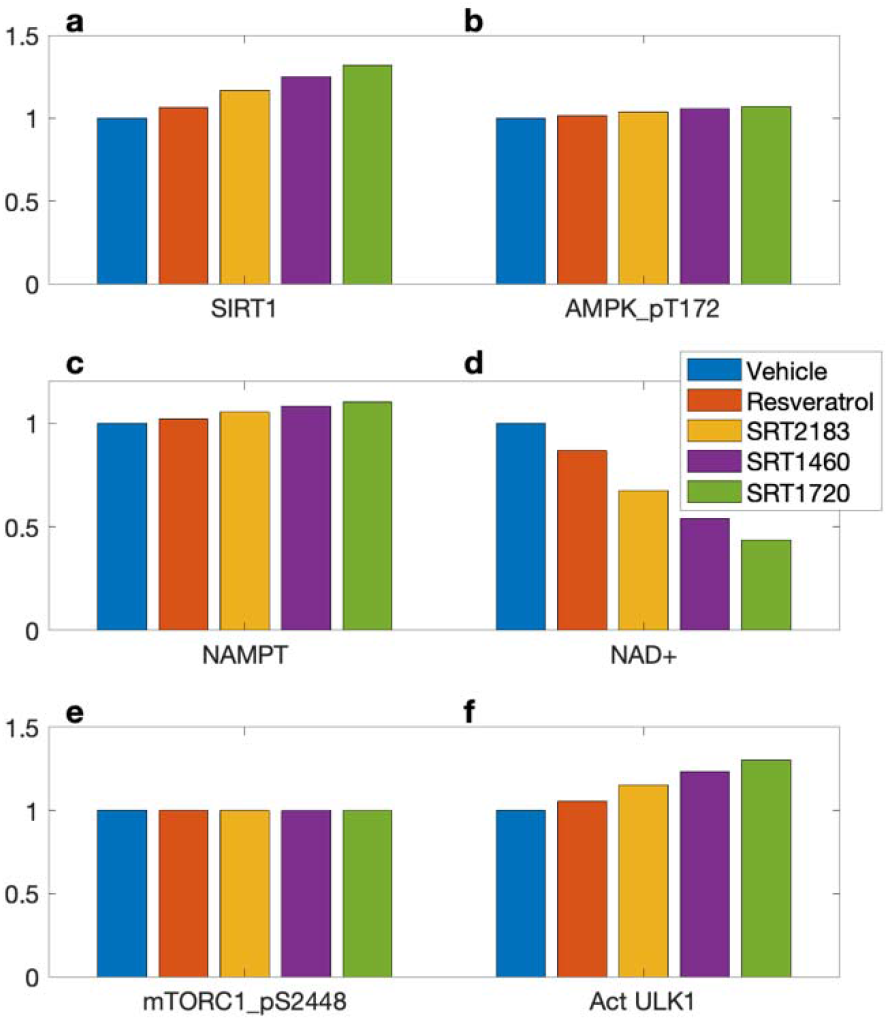
Predicted effects of STACs on SIRT1 activity (panel ***a***), AMPK_pT172 (***b***), NAMPT (***c***), NAD+ (***d***), mTORC1_pS2448 (***e***), and activated ULK1 (***f***). Values are shown relative to control.

AMPK regulates nicotinamide phosphoribosyl transferase (NAMPT),(27) which acts as a limiting enzyme in the conversion of NAM to nicotinamide mononucleotide (NMN). Taken in isolation, the STAC-induced elevation in NAMPT (Fig. 7c) would accelerate the conversion of nicotinamide (NAM) to NMN, resulting in an increase in the downstream NAD+. However, the STACs also accelerate the consumption of NAD+ by binding on the allosteric site of SIRT1. Indeed, the enhanced SIRT1 consumption is the stronger effect, resulting in lower NAD+ following the administration of STACs (Fig. 7d), by as much as 56% with SRT1720.

The present model is based on C2C12 cells, in which AMPK does not regulate mTORC1 directly but instead facilitates the phosphorylation of TSC1_TSC2 at ser S1387.(9) Because both TSC1_TSC2 and its phosphorylated form TSC1_TSC2_pS1387 deactivate mTORC1, AMPK has a negligible effect on mTORC1 inhibition. AMPK regulates autophagy through direct phosphorylation of ULK1.(21) Thus, despite mTORC1’s insensitivity to STACs, the level of activated ULK1 is predicted to be increased by STACs (Fig. 7f).

### STACs may induce premature autophagy

The inhibitory phosphorylation of ULK1 by mTORC1 serves as a negative feedback to prevent overactive autophagy and to avoid a futile cycle in which newly synthesized cellular building blocks are prematurely broken down again. Given that STACs activate ULK1, we investigate the possibility that STAC intake may lead to premature autophagy, i.e., protein catabolism under an abundance of nutrients. We consider three groups: control, SRT1720, and a (hypothetical) higher-performing STAC (called “SRTx”) which we assume reduces substrate K_m_ by 90%. Simulations are conducted for baseline AA level, and a low AA level that is 25% of baseline. Predicted AMPK_pT172, mTORC1_pS2448, and ULK1 activation levels are shown in Fig. 8. Simulations predict that, with abundance of nutrients, SRTx yields ULK1 activation level that is 91% of the analogous level obtained for the control group under AA deprivation. The similar autophagy activation under these two sets of conditions, despite the widely differing AA abundance levels, suggests that a sufficiently potent STAC may induce premature autophagy.

**Figure 8.**
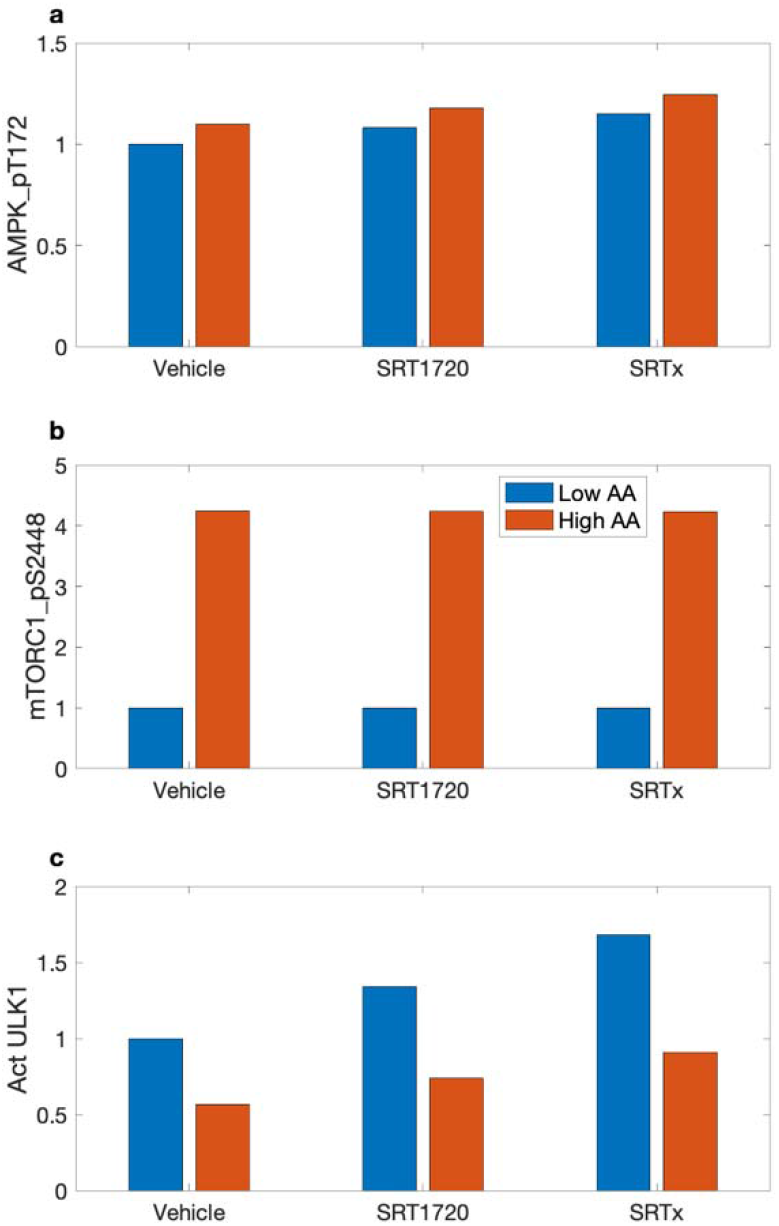
Potential overactivation of autophagy by STACs at high nutrient level. ***a***, normalized levels of AMPK_pT172 obtained for control (“Vehicle”), for SRT1720, where SIRT1 substrate K_m_ is reduced by 70%, and for a hypothetical STAC where K_m_ is reduced by 90% (“SRTx”). For each case, AMPK_pT172 levels are determined at an AA level is 25% of control (“Low AA”) and for reference AA conditions (“High AA”). ***b***, normalized levels of mTORC1_pS2448. ***c***, activated ULK1. With a sufficiently strong STAC (“SRTx” and “High AA”), autophagy may be activated at a level close to that at AA deprivation (compare with “Vehicle” and “Low AA”).

### Parameter sensitivity analysis

We perform local sensitivity analysis to assess the response of model outputs to small variations in selected parameter values (Fig. 9). Two distinguishable heat-map regions are identified with mostly non-zero entries. These regions are associated with the original model components: the insulin/IGF-1 signaling pathway (top-left region) and the Preiss-Handler and salvage signaling pathways (bottom-right region). These two components are coupled via the effects of AMPK on NAMPT, and of SIRT on AMPK. Variations in parameters in one region have typically minor (but nonzero) effects on outputs in the other region.

**Figure 9.**
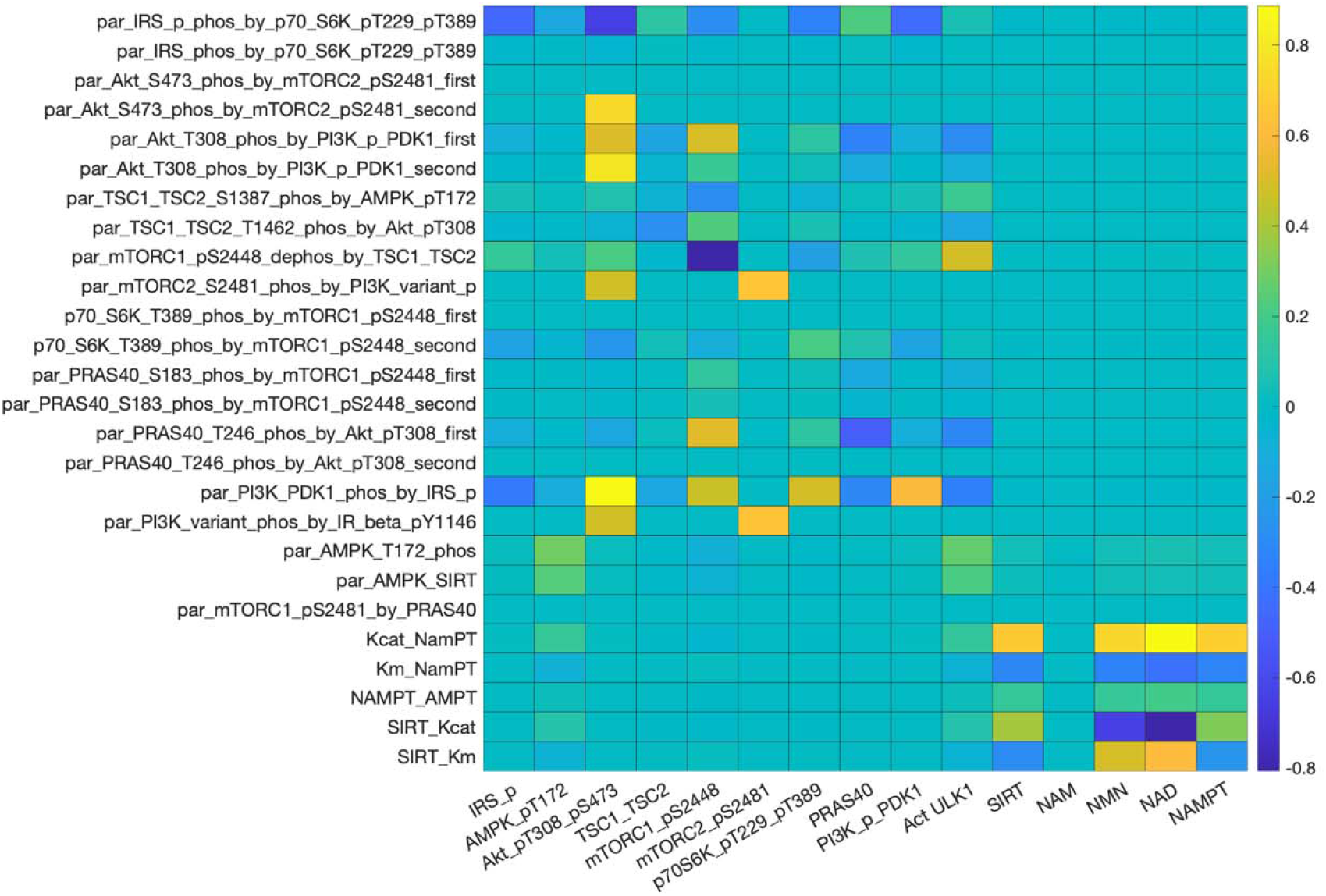
Heat map that illustrates the sensitivity of key model outputs (horizontal axis) to local variations in selected model parameters (vertical axis). Definition of the parameters can be found in the Supplemental Materials.

Within the insulin/IGF-1 signaling pathway, model outputs are particularly sensitive to variations in links that activate or inhibit proteins that have a large number of downstream effects. E.g., the activation of IRS_p by p70S6K_pT229_pT389 (“par_IRS_phos_by_p70_S6K_pT229_pT389”). Except for mTORC2_pS2481, other outputs in the insulin/IGF-1 signaling pathway exhibit significant sensitivity to variations in that activation strength, because IRS_p directly modulates major model variables such as PI3K_PDK1 and AMPK. In contrast, model solution is insensitive to changes in the activation of IRS_pS636 by p70S6K_pT229_pT389, in large part because the IRS_pS636 has no direct downstream effect.

Within the Preiss-Handler and salvage signaling pathways, most model outputs exhibit significant sensitivity to variations in the selected parameters associated with the pathways. NAM exhibits remarkable stability, because of the robust negative feedback cycle involves itself, NMN, NAD+, and SIRT1. Proteins in this pathway are particularly sensitive to changes in the abundance of active NAMPT (“Kcat_NAMPT”). This result suggests that an NAMPT activator may be a promising pharmacological approach for raising intracellular NAD+ and SIRT1, to realize diverse and potentially impactful therapeutic benefits.

## DISCUSSION

Traditionally, the investigation of human ageing and disease has relied on cell cultures and animal models, including non-vertebrates (e.g., yeast, worm, and fly) and vertebrates (e.g., zebrafish, mice, dogs, and primates), as well as clinical trials. With the advent of bioinformatics and computational biology, computational modeling and analysis techniques can provide accurate simulation of biological processes. The present study describes a state-of-the-art computational model for investigating signaling pathways in growth, ageing, metabolism, and disease in mammals. Major model components include the insulin/IGF-1 or mTOR signaling pathway,(28) the Preiss-Handler and salvage pathways,(29) energy sensor AMPK, and transcription factors FOXO and PGC-1α. The mTOR signaling pathway couples energy and nutrient abundance to the execution of cell growth and division. That function can be attributed to the ability of mTORC1 to sense energy, nutrients, and growth factors, by regulating other important kinases, such as S6K and AKT. The Preiss-Handler and salvage pathways regulate the metabolism of NAD+ as well as NAD+-consuming proteins such as sirtuins. Key findings include:

- Model simulations indicate that simultaneously inhibiting AKT or PI3K_PDK1, and inhibiting mTORC1 effectively suppresses PRAS40 phosphorylation on both Ser183 and Thr246 sites, further enhancing the inhibition of mTORC1 by PRAS40 (Fig. 1). This result suggests a clinically important role of PRAS40 in controlling tumor growth.
- We provide the first computational model that capitulates the interplay between sestrin2 and leucine, and the effect on mTORC1 activity. With a crucial role in metabolic regulation through the activation of AMPK and inhibition of mTORC1, sestrin2 might serve as a therapeutic target for cancers, metabolic diseases, and neurodegenerative diseases.
- Given that sestrin2’s inhibitory effect on mTORC1 activity can be significantly impacted by AAs such as leucine (Fig. 5), dietary modifications may enhance the efficacy of therapies that target mTORC1 and sestrin2.
- The model captures the interactions between arginine and leucine during protein deprivation, and predicts a signal that reactivates mTORC1 and downregulates autophagy (Fig. 4).
- The model capitulates the regulation of autophagy by SIRT1. Simulations indicate that, by activating SIRT1, STACs in high dosages may lead to premature autophagy (Fig. 8).

mTORC1 signaling is switched on by a number of oncogenic signaling pathways and may be hyperactive in up to 70% of all human tumors.(30) Thus, there is much interest in targeting mTORC1 signaling as a potential therapeutic avenue for anti-cancer therapy. Rapamycin, originally developed as an immunosuppressant that targets T-cells, is arguably the best known mTORC1 inhibitor and has been shown to extend lifespan in mice.(31) However, despite its specificity, rapamycin in typical dosages does not completely inhibit all mTORC1 activities,(32) limiting its efficacy as an anti-cancer agent. In fact, cancer patients whose tumors exhibit a mutational activation of PI3K/AKT signaling have a low response rate for rapamycin and its rapalogs (e.g. breast, colon and prostate cancer, and glioblastoma(33, 34)). This inadequate therapeutic response is believed to result from rapamycin and its rapalogs’ incomplete inhibition of mTORC1-mediated phosphorylation of 4E-BP1 and a concomitant activation of AKT via loss of a negative feedback mechanism.(28)

Mi et al. suggested that the acquired resistance to rapamycin in cancer cells may be attributable to the redundant phosphorylation of PRAS40 by both AKT and mTORC1 signaling.(35) Their findings indicate that the concurrent inhibition of AKT and mTORC1 yields effective inhibition of PRAS40 phosphorylation on both Ser183 and Thr246 sites, enhancing PRAS40’s inhibition of mTORC1-mediated 4E-BP1 phosphorylation and translation, concomitant with suppression of tumor growth and cell motility.(35) Similarly, while PI3K inhibitors such as wortmannin are potential anti-cancer agents on their own,(36) a more nearly complete inhibition of mTORC1 may be achieved when inhibitors of the PI3K/AKT pathway are administered in conjunction with rapamycin (Fig. 1). These results suggest a potentially important role of PRAS40 in the translational control of tumor progression. Indeed, dual inhibition of PI3K and mTORC1 signaling by rapalogs in combination with PI3K or AKT inhibitors has demonstrated profound efficacy in preclinical cancer models.(37–40)

Chronic administration of rapamycin leads to insulin resistance due to its suppression of mTORC2, resulting in glucose intolerance and hepatic insulin resistance.(19) The loss of insulin sensitivity is due to the reduction of AKT_T308_S473, which plays a pivotal role in the translocation of GLUT4.(14, 15) Sebastian and co-workers demonstrated intermittent administration of rapamycin (e.g., once every 5 days) mitigates its detrimental effect on glucose homeostasis.(16, 18) Our model simulations indicate that an optimal rapamycin dosage can be identified which, in chronic and continuous usage, attenuates the detrimental effect on insulin sensitivity while preserving rapanycin’s anti-cancer and anti-ageing effects via mTORC1 inhibition (Fig. 3).

Sestrin2 is a highly conserved stress-inducible metabolic protein known to provide protection to cells against oxidative stress, endoplasmic reticulum (ER) stress, and hypoxia. Sestrin2 also plays a key role in metabolic regulation through the activation of AMPK and inhibition of mTORC1, with downstream effects including autophagy activation, antiapoptotic effects in normal cells, and proapoptotic effects in cancer cells.(41) As such, sestrin2 might serve as a potential therapeutic target for cancers,(42),(43) metabolic diseases, and neurodegenerative diseases.(44)

The present model provides the first computational platform that capitulates the regulation of mTORC1 by sestrin2, and its modulation of leucine. The action of sestrin2 on mTORC1 begins with its inhibition of GATOR2. During acute starvation, sestrin2 binds to GATOR2 and impedes the latter’s inhibition of GATOR1, resulting in the suppression of mTORC1. Sestrin2 binds with leucine; the addition of leucine would reduce free sestrin2, promote the downstream inhibition of GATOR1 and ultimately enhance the activation of mTORC1. The interplay between sestrin2 and leucine, and the effect on mTORC1 activity is captured in Fig. 5. As noted above, sestrin2 is a potential target for anti-cancer and other disease treatments. Given that sestrin2’s inhibitory effect on mTORC1 activity can be significantly impacted by AAs such as leucine, dietary modifications may enhance the efficacy of therapies that target mTORC1 and sestrin2.

GATOR1 negatively regulates mTORC1 and appears to function as a tumor suppressor. In cancer cells with loss-of-function mutations in GATOR1, which include some lung, breast, ovarian cancers, and glioblastomas,(45) mTORC1 is hyperactive and these cancer cells have been shown to be hypersensitive to rapamycin. Thus, the associated cancers may be particularly amenable to therapeutic strategies that limit mTORC1 activity. For rapamycin-resistant cancers, and for diabetes and neurodegenerative diseases, GATOR may be a potential therapeutic target. In fact, as an upstream regulator of mTORC1, GATOR1 has been identified as a potential therapeutic target for epilepsy, in which mTORC1 hyperactivation has been implicated.(46)

Also contributing to cellular AA sensing for the mTORC1 pathway is the CASTOR family.(47) CASTOR1/2 binds to GATOR1 and obstructs the binding process between GATOR1 and GATOR2, freeing up GATOR1 to inhibit mTORC1. Unlike sestrin2 which is inhibited by leucine, CASTOR1 binds to arginine. Thus, the addition of arginine elevates mTORC1 activity by reducing free CASTOR1.

Another arginine sensor is the lysosomal AA transporter SLC38A9. Arginine stimulates SLC38A9 and promotes its interaction with the Rag GTPase-Regulator complex, which results in the activation of mTORC1. Wyant et al. reported that a mutant of SLC38A9 that does not interact with arginine lacks the ability to signal AA sufficiency to mTORC1.(48) Another important role of SLC38A9 in AA homeostasis is to transport AAs produced by lysosomal proteolysis, such as leucine, from lysosomes to the cytosol, thereby reactivating mTORC1. The critical role of SLC38A9 in this process is evinced by the SLC38A9-null HEK293T cells, which exhibit whole-cell AA levels similar to wild-type, but significantly higher lysosomal AA concentrations including leucine.(48) The efflux of lysosomal leucine and the subsequent activation of mTORC1 may regulate autophagy. During starvation, mTORC1 is inhibited, which attenuates its inhibitory phosphorylation of ULK1 and promotes autophagy. Thus, intracellular nutrients produced by autophagy can stimulate mTORC1 signaling and provide a negative feedback signal to downregulate autophagy (more below).(20) The present model is unique in that it explicitly and separately represents arginine and leucine, rather than lumping them into a single AA category. Consequently, the model captures the exquisite interactions of these AAs, and the resulting signal to the mTORC1 pathway and its reactivation during protein deprivation. Indeed, the model is the first to recapitulate the reactivation of mTORC1 by proteolysis following autophagy (Fig. 4).(20)

STACs enhance autophagy via its activation of SIRT1, followed by the activation of AMPK and inhibition of the mTORC1. The elevated SIRT1 activity also facilitates the deacetylation of FOXOs, which are transcription factors that have major impacts on longevity and cancer. In the absence of growth factors or AAs, the reduction in activated AKT leads to the dephosphorylation of FOXOs and drives their relocalization from the cytoplasm to the nucleus. The subsequent deacetylation of FOXOs by SIRT1 mediates stress resistance response. AMPK phosphorylation also increases the transcription activity of FOXOs. Emerging evidence suggests that FOXO factors act as a tumor suppressor in a variety of cancers.(49)

Additionally, SIRT1 regulates autophagy through the direct deacetylation of autophagy-related genes such as ATG. Studies have demonstrated an essential role for SIRT1 in the induction of autophagy(50, 51), as well as the protective effects of the SIRT1-induced autophagy in preventing or attenuating neurotoxicity.(52, 53) However, by activating SIRT1, STACs may lead to the decoupling of the autophagic response from the organisms’ nutrient and energy status. Model simulations suggest that a sufficiently potent STAC may yield premature autophagy (Fig. 8), in which newly synthesized cellular building blocks are prematurely and unnecessarily catabolized despite an abundance of nutrients.

The insulin/IGF-1 signaling pathway of the present model is fitted primarily for C2C12 cells,(10) whereas the Preiss-Holder and salvage pathway is formulated primarily using nonspecific mammalian cell data.(12) Because different cells have diverse energetic and metabolic needs, their metabolic pathways often exhibit significantly different functions and regulations. For instance, SIRT1 is activated by starvation in most cells,(54) but in the mouse pancreas, SIRT1 is inactivated by the reduction in the NAD+/NADH ratio.(55) In C2C12 cells, AMPK does not directly regulate mTORC1 but promotes TSC1_TSC2 phosphorylation at ser S1387. Because mTORC1 is inhibited by both TSC1_TSC2 and its phosphorylated form TSC1_TSC2_pS1387, in C2C12 cells AMPK appears to have only a limited regulatory effect on mTORC1. In contrast, AMPK is known to directly inhibit mTORC1 in HEK293T cells.(56) Furthermore, sex and age differences have been reported. In the mouse, mTORC1 and mTORC2 activity levels are known to vary between males and females.(57) In human skeletal muscle, mTORC1 is activated under different conditions depending on the age of the individuals.(58, 59) Similarly, marked changes have been reported in SIRT1 activity and NAD+ levels as individuals age.(60, 61) These observations highlight the importance of taking into account cell or tissue specificity, together with sex and age, when developing therapeutic strategies that target these pathways. Investigating how the key ageing regulatory pathways interact during normal ageing is a worthwhile future extension.

In sum, we have developed a state-of-the-art computational model for investigating the interactions among signaling pathways and environmental stimuli in growth, ageing, metabolism, and diseases. The model can be used as an essential component to simulate (1) gene manipulation, (2) therapies for cancer, metabolic diseases, and neurodegenerative diseases, (3) calorie restrictions, and (4) chronic stress. Simulation results can be interpreted to assess the implications on longevity and ageing□related diseases.

## MATERIALS AND METHODS

Main model components include the insulin/IGF-1 or mTOR signaling pathway(28), the Preiss-Handler and salvage pathways,(29) energy sensor AMPK, and transcription factor FOXO and PGC-1α, in the mouse C1C12 cell. The dynamics of the signaling pathways is modeled as a system of ODEs; Figs. 10 and S1 depict the pathways and protein interactions. The reactions and associated parameters are presented in Table S1 and the Excel file. Model parameters are taken from published studies (10, 12) and fitted to experimental data (62, 63) using the interior point optimization method.

**Figure 10.**
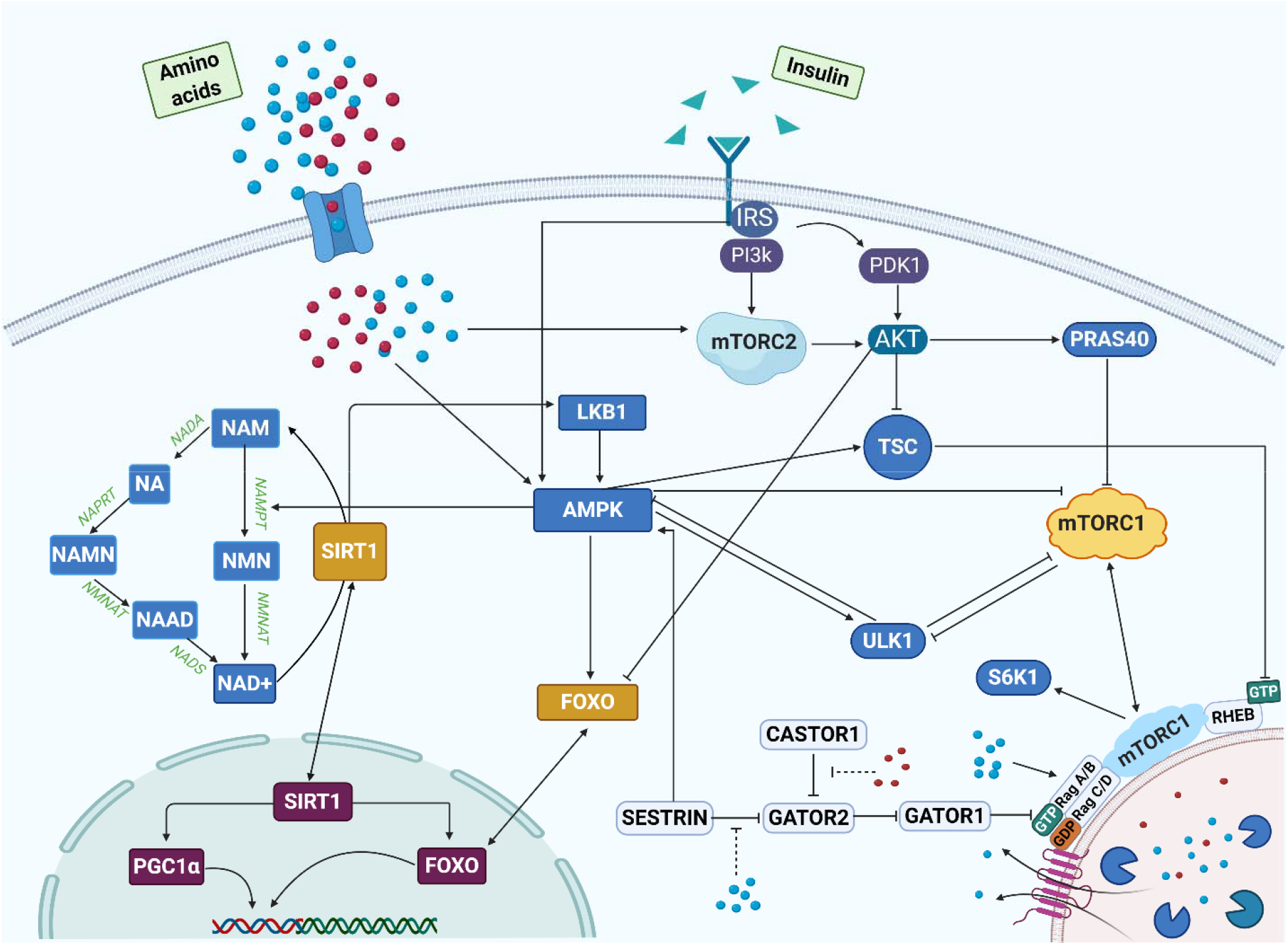
Pathway representation of network model, some components of which may be activated by amino acids, such as leucine (blue circles) and arginine (red circles), and insulin (green triangles0. The model represents three distinct areas in the cell: the cytoplasm, lysosome, and the nucleus.

Within the insulin/IGF-1 signaling and mTOR pathways, insulin activates the insulin receptor (IR), which triggers the IRS and PI3K, resulting in the phosphorylation of PDK1 and mTORC2, respectively. mTORC2 phosphorylates AKT on the S473 and T308 residues, whereas PDK1 activates AKT. Active AKT phosphorylates a variety of proteins, including FOXOs, TSC1_TSC2 and PRAS40. Phosphorylation of TSC1_TSC2 by AKT inactivates the TSC complex, thereby activating mTORC1 and resulting in a number of downstream effects. These include (i) inhibition of ULK1 by phosphorylating ULK1 on Ser 757; the ULK1 complex is a key contributor to the initiation of autophagy; (ii) phosphorylation of PRAS40, which attenuates the inhibition of mTORC1; (iii) phosphorylation of p70S6K on T229 and T389, which initiates a negative feedback loop via IRS.

An important novel aspect of our model is the explicit representation of the role of sestrin2 and leucine in the regulation of mTORC1. Sestrin2 deactivates mTORC1, via its effects on protein complexes GATOR1 and GATOR2. GATOR1 inhibits mTORC1, whereas GATOR2 inhibits GATOR1. Sestrin2 inhibits GATOR2, enhancing the activation of GATOR1, and eventually suppressing mTORC1. This process is modulated by leucine, which binds to sestrin2 and impeding its inhibitory effect on mTORC1. Thus, mTORC1 activation depends, among other factors, on the concentration of sestrin2 and leucine. To illustrate that dependence, we consider the mTORC1 activation rate:

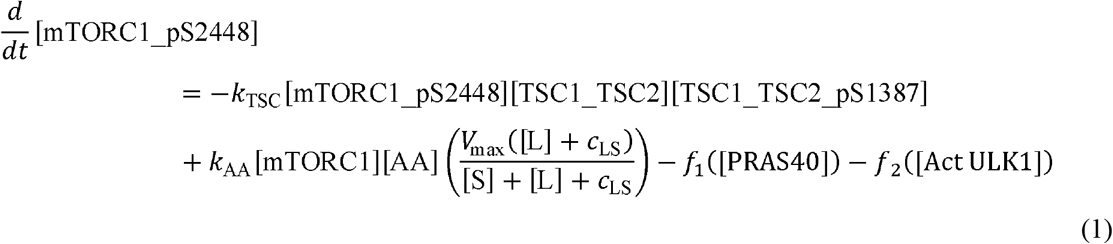

where [A], [L], and [S] denote the arginine, leucine, and sestrin2 concentrations, respectively; and mTORC1_tot_ denotes total mTORC1. Act ULK1 denotes the activated form of ULK1. The first term on the right describes the inhibition of mTORC1 by the TSC1_TSC2 complexes; the second term describes the effects of leucine and sestrin2 on mTORC1; the third and fourth terms describe the contributions from PRAS40 and activated ULK1.

The model simulates the dynamics of another key player in metabolism, NAD+, which is produced through the *de novo*, Preiss-Handler and salvage pathways. The major source of NAD+ in mammals is the salvage pathway, which recycles NAM produced by enzymes utilizing NAD+. The first step in the salvage pathway involves the rate-limiting enzyme NAMPT, which facilitates the conversion of NAM to NMN and whose activity is increased by AMPK, and the expression and/or activity of which when reduced is associated with ageing and poor health. (64) The second step converts NMN to NAD+ via the NMNAT enzymatic reaction. NAD+ thus produced activates substrates including SIRT1 and is consumed in the process. Albeit less important for NAD+ biosynthesis in mammals, the Preiss-Handler pathway is also represented. The pathway begins with the conversion of NAM to NA, followed by the conversion to NAMN, catalyzed respectively by NADA and NAPRT. Like the salvage pathway, NMNAT in Preiss-Handler pathway also catalyzes the process of production of NAAD from NAMN. Finally, the reamidation of NAAD by NADS yields NAD+. The Preiss-Handler pathway is more energy and ATP consuming than the salvage pathway.

Connecting the mTORC1 and NAD+ pathways is AMPK, a master regulator of cellular energy that is activated under starvation or hypoxia. AMPK can be stimulated by sestrin, IRS, and liver kinase B1 (LKB1) which deacetylated by SIRT1. Activated AMPK promotes autophagy by directly phosphorylating and activating ULK1. As such, there is a competition between mTORC1 and AMPK to phosphorylate different residues of ULK1 to decide cells fate. ULK1 in turn inhibits AMPK and mTORC1 in a negative feedback loop, whereas leucine activates them.

Therapeutic treatments and dietary conditions are simulated by changing selected model parameters (see below). For a given set of parameters, a steady-state solution can be computed by integrated the model equations for a sufficiently long time. For simulations that involve a change in model parameters, model variables are initialized to the steady-state solution corresponding to the parameter values at the initial time. Such initial conditions are realistic and avoid an abrupt change in solution at the initial time.

### Simulating the administration of rapamycin and wortmannin

To simulate acute administration of rapamycin, we decrease total mTORC1 by 75%.(17) To simulate chronic administration of rapamycin, we decrease the total amount of both mTORC1 and mTORC2 by 75%.(17) To simulate the effect of wortmannin, we reduce total PI3K_PDK1 by 80%.(65)

### Simulating the glucose tolerance test

Chronic administration of rapamycin is known to inhibit mTORC2 and impair one’s insulin sensitivity by reducing AKT_pT308_pS473, which is essential in the translocation of GLUT4.(14, 15) To determine an optimal rapamycin dosage that attenuates its detrimental effect on insulin sensitivity while preserving its inhibition of mTORC1, we simulate the glucose tolerance test and track plasma glucose concentration by

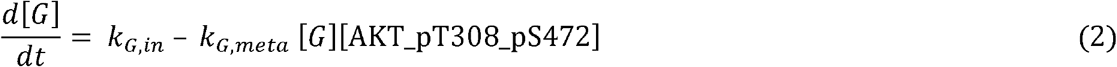

The cellular uptake rate of plasma glucose is assumed to be proportional to the cellular concentration of AKT_pT308_pS472. As in the previous simulations, the model is initialized at the fasting state with a low insulin level, glucose = 1, and *k_G,in_* = 0 which represents the absence of glucose intake; additionally, *k_G,meta_* = 7×10^−4^/min. At t = 40 min, the glucose tolerance test begins, whereby plasma glucose is elevated by setting *k_G,in_* to 0.115/min at t = 40 min then linearly decreases to 0 in the next 25 minutes. Additionally, insulin is proportional to plasma glucose.

### Simulating the regulation of mTORC1 by arginine and leucine

SLC38A9, a lysosomal 11-transmembrane-segment protein with homology to AA transporters, is needed to transport, in an arginine-regulated fashion, most essential AAs out of lysosomes, including leucine.(66) Wyant et al. reported that arginine enhances the capacity of SLC38A9 to transport leucine, by increasing its V_max_ without significantly affecting its K_m_.(48) Taking both leucine and arginine into account, we model leucine transport rate, denoted V_max,L_, as

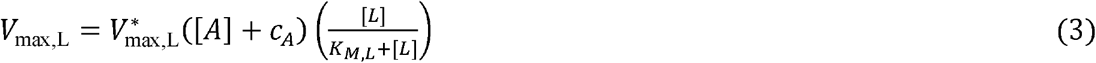

where c_A_ denotes the contribution of lysine, which also transports leucine albeit significantly less effectively than arginine. In the absence of either AA groups ([A] + c_A_ = 0 or [L] = 0), there is no leucine efflux. Under basal conditions, the concentration of lysine is sufficiently low that c_A_ is taken to 0.

### Simulating the regulation of mTORC1 by sestrin2 and leucine

Sestrin2 and leucine exert opposite effect on mTORC1, with sestrin2 deactivating mTORC1, and leucine increasing mTORC1 activity by binding to sestrin2. To investigate the dependence of mTORC1 activation level on leucine and sestrin2, we consider the steady-state formulation of Eq. 1 (by setting the time-derivative to zero) and solve for [mTORC1_pS2448]. We obtain

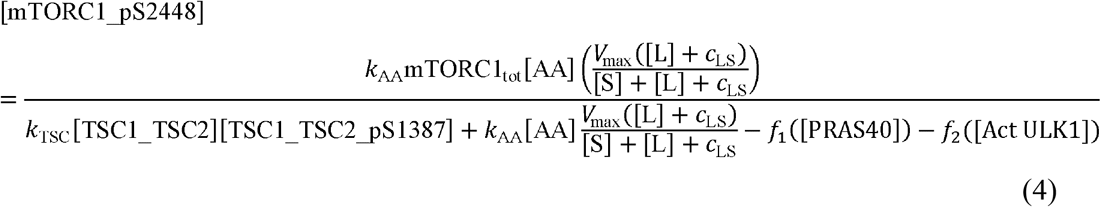

### Simulating STACs and their anti-ageing effects

To illustrate the effects of resveratrol and other STACs, we assume that the dependence of SIRT activity on NAD+ can be approximated by Michaelis-Menten kinetics.(67, 68) Based on in vitro findings by Milne et al. at 10 μM, resveratrol reduces substrate K_m_ by 20%, SRT2183 by 50%, SRT1460 by 60%, and SRT1720 by 70% (figure 2a in Ref. (69)). As shown in Fig. 6a, at baseline K_m_ of [NAD+] = 0.029 mM, resveratrol slightly enhances SIRT1 activity by 11%. Other STACs has larger effects; SRT2183 elevates SIRT activity by 29%, SRT1460 by 43%, and SRT1720 by 54%.

### Parameter sensitivity analysis

Sensitivity of model output *x* to a parameter *p* is given by the relative change in *x* with respect to a 1% relative change in *p*, i.e.,

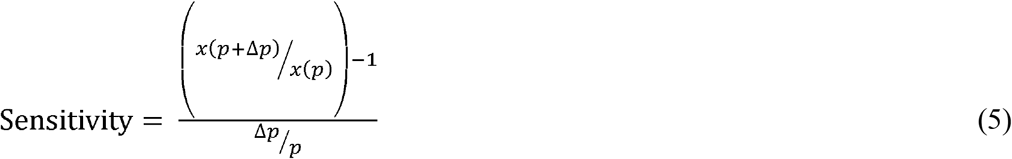

where in our computations, Δ*p* = 0.01 *p*. Other parameters are fixed and all model outputs are updated simultaneously when *p* is varied.

### Data availability

MATLAB programs used in the model simulations can be accessed at https://github.com/MehrshadSD

## Acknowledgements

This research was supported by the Canada 150 Research Chair program and the NSERC Discovery award.

## Support information

**Figure S1.** Schematic diagram depicting connections among model components that form the signaling pathways.

**Figure S2.** Effects of insulin, rapamycin, and wortmannin on key proteins and their interactions.

**Figure S3.** Effect of protein deprivation and subsequent leucine efflux on key model variables, obtained for differing arginine levels.

**Figure S4.** Effect of protein depletion and restoration on key model variables, obtained for differing sestrin2 levels.

## Model equations

The model and parameters that support the findings of this study are available from the corresponding author.

