## Supplementary information 1 for "Interactions among mTORC, AMPK, and SIRT: A Computational Model for Cell Energy Balance and Metabolism"

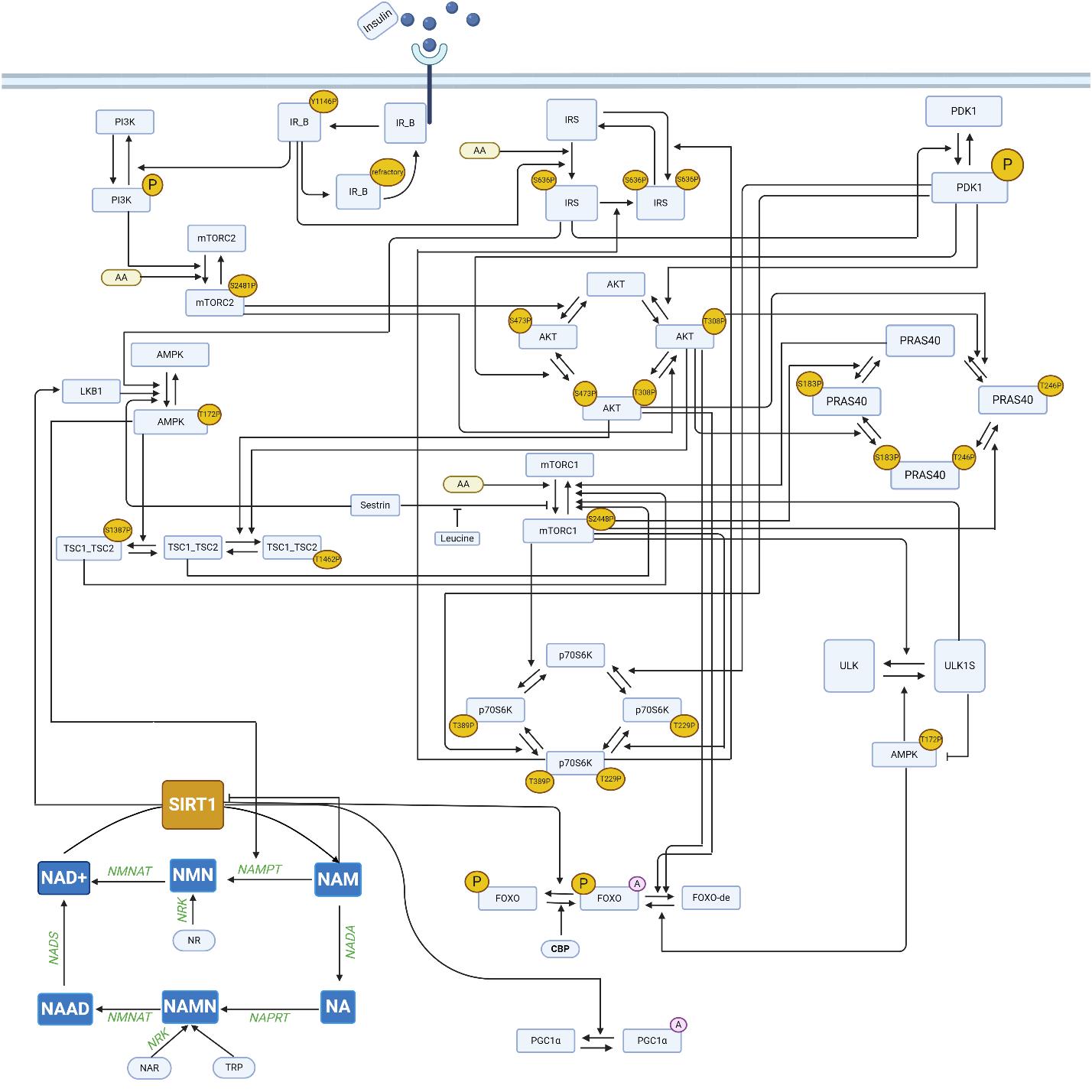

**Figure S1.** Schematic diagram depicting connections among model components that form the signaling pathways. AA represents amino acids; “A” in purple circles denote the acetylated form of the complex; “P” in yellow circles denote the phosphorylated form.

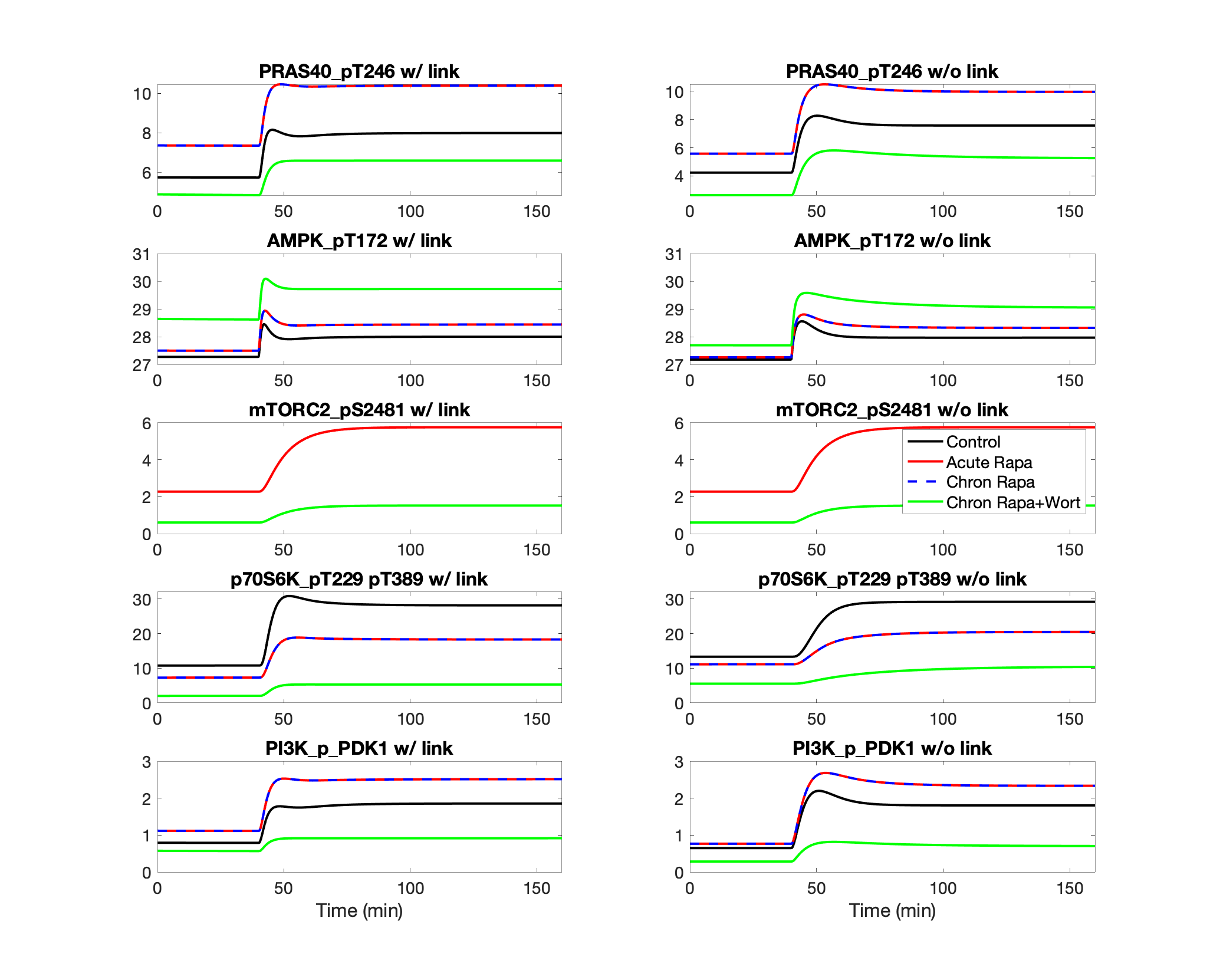

**Figure S2.** Effects of insulin, rapamycin, and wortmannin on key proteins and their interactions. Insulin was lowered to 10% of its baseline level for the initial 40 min of the simulation, and subsequently returned to baseline level. Simulations are conducted for control, acute and chronic administration of rapamycin, chronic administration of rapamycin with wortmannin. The inhibition of PRAS40 of mTORC1 is represented in the left panels but not the right ones. Model predicts that PRAS40 substantially lowers mTORC1 level under low insulin conditions. That effect is the most prominent under chronic administration of rapamycin and wortmannin.

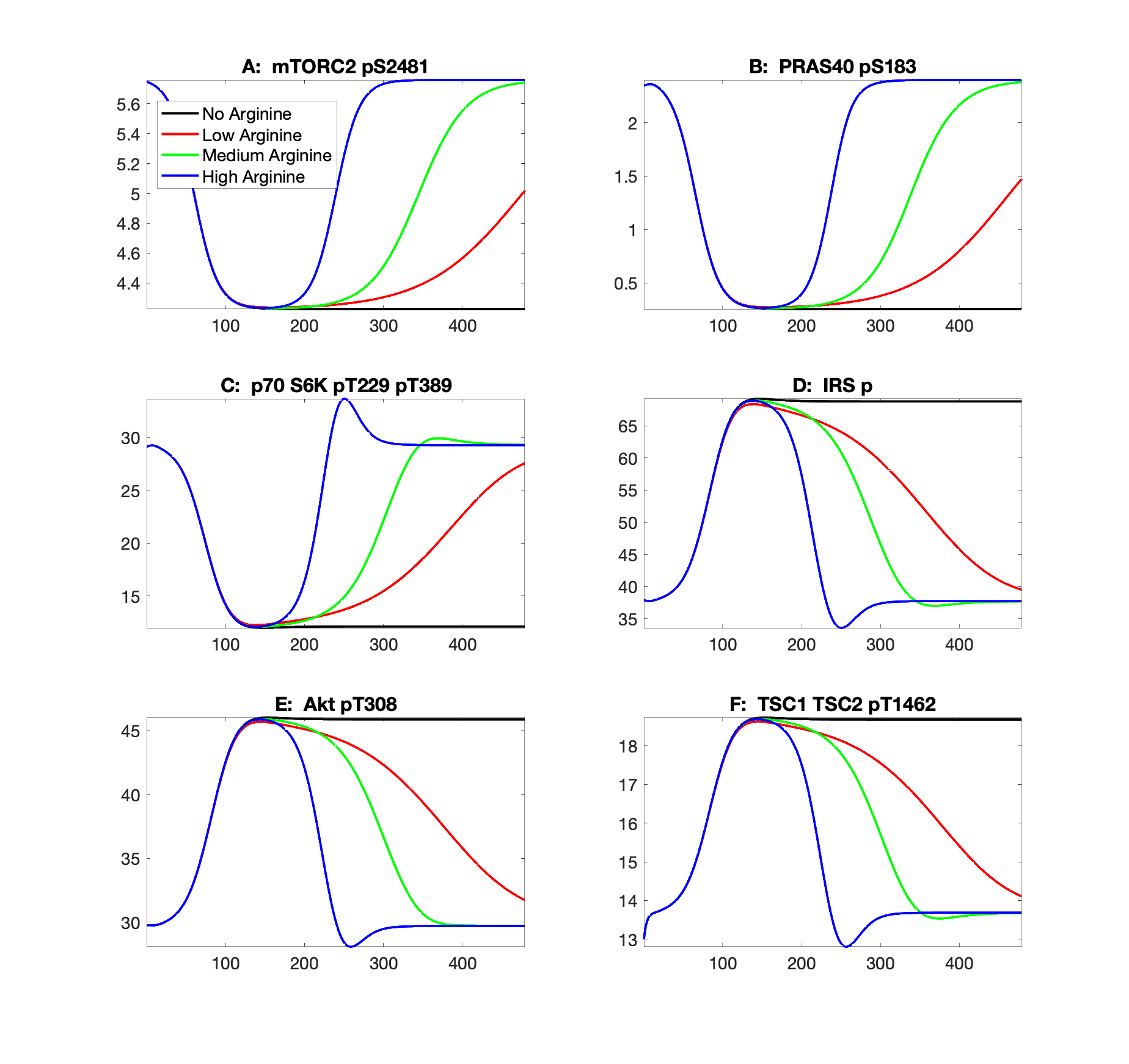

**Figure S3.** Effect of protein deprivation and subsequent leucine efflux on key model variables, obtained for differing arginine levels.

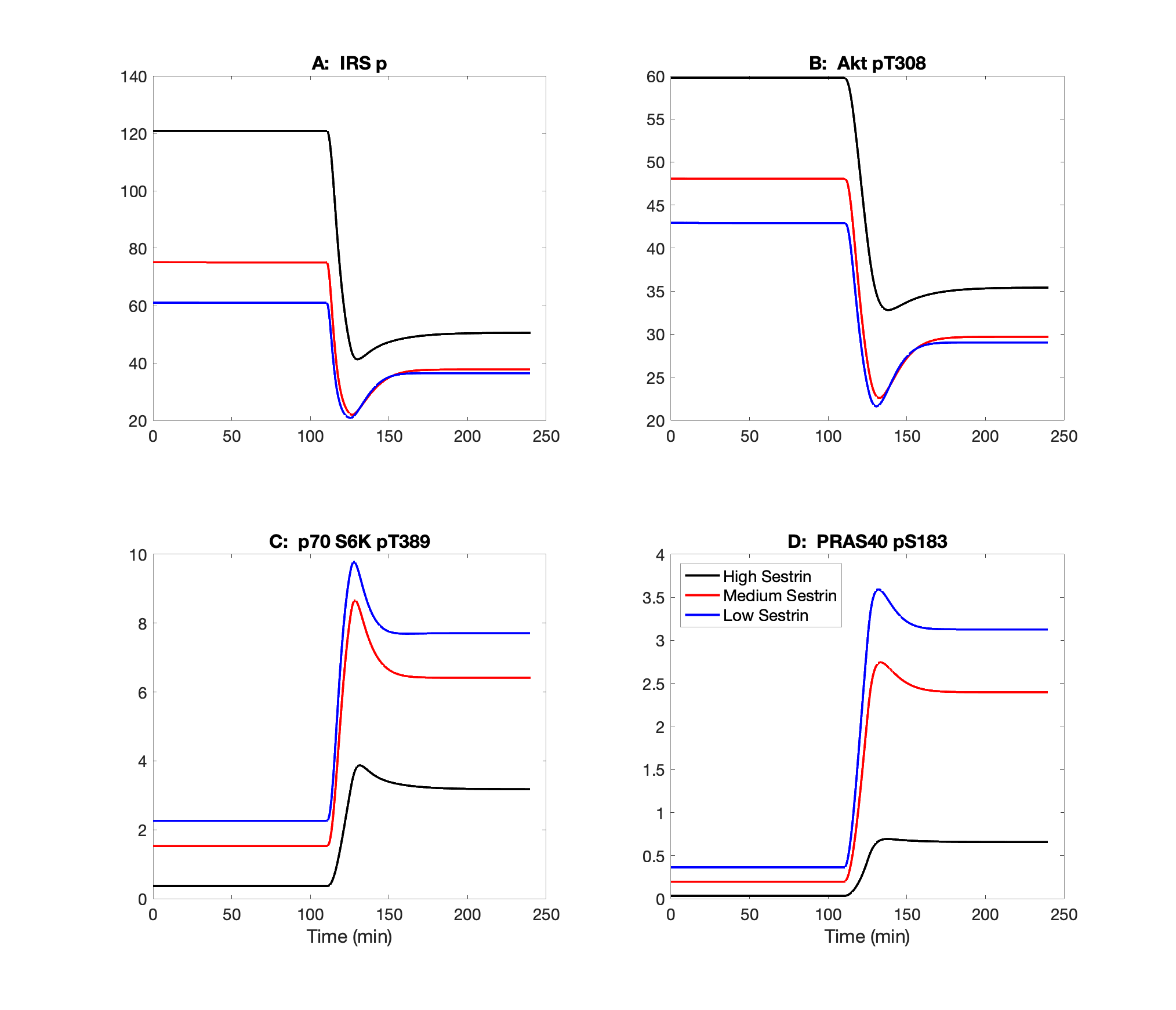
 **Figure S4.** Effect of protein depletion and restoration on key model variables, obtained for differing sestrin2 levels.
